# CMA-ES-Rosetta: Blackbox optimization algorithm traverses rugged peptide docking energy landscapes

**DOI:** 10.1101/2022.12.19.521113

**Authors:** Erin R. Claussen, P. Douglas Renfrew, Christian L. Müller, Kevin Drew

## Abstract

Energy minimization is necessary for virtually all modeling and design tasks and involves traversing extremely rugged energy landscapes. Although the gradient descent based minimization routines in Rosetta have fast runtimes, due to these rugged landscapes, minimization often converges into high-energy local minima. Alternative numerical optimization techniques, such as evolution strategies, are more robust to rugged landscapes and have been shown to be highly successful on a diverse set of problems. Here we explore the Covariance Matrix Adaptation Evolution Strategy (CMA-ES), a state-of-the-art derivative-free optimization algorithm, as a complementary approach to the default minimizer in Rosetta. We used a benchmark of 26 peptides, from the FlexPepDock Benchmark, to assess the performance of three algorithms in Rosetta, specifically, CMA-ES, Rosetta’s default minimizer, and a Monte Carlo protocol of small backbone perturbations. We test the algorithms’ performance on their ability to dock an idealized peptide to a series of hotspots residues (i.e. constraints) along a native peptide. Of the three methods, CMA-ES was able to find the lowest energy conformation for 23 out of 26 benchmark peptides. The application of CMA-ES allows for an alternative optimization method for macromolecular modeling problems with rough energy landscapes.

## Introduction

The computational modeling of biomolecular structure is invaluable for determining insights into molecular mechanisms of biomolecules. As the field of macromolecular modeling has expanded into a vast array of applications from protein structure prediction^1^ to small molecule docking^2^ to protein design^3^ and more, new protocols have been developed in order to accommodate developers’ needs^4^. Although different strategies have been devised, nearly all modeling protocols rely on the extremely fast class of gradient descent energy minimization algorithms to find the lowest energy conformation^5^.

Current energy minimization algorithms, although fast, often get stuck in local energy minima due to high energy non-native conformations in the energy landscape^6^. Energy functions that estimate the energy landscapes of macromolecules range from coarse grain which are smooth but inaccurate to high-resolution which are more accurate but also more rugged (i.e. have many local minima)^7^. Present day modeling and design applications require accurate high-resolution energy functions. Consequently, there is a need for new minimization techniques better suited for dealing with rougher energy landscapes and their inherent local energy traps.

Alternatives to gradient descent minimization, such as evolutionary optimization strategies, are attractive because they are more robust to rough gradients due to their ability to escape local minima^8^. One such algorithm, covariance matrix adaptation evolutionary strategy (CMA-ES), is a stochastic gradient-free method for optimizing non-convex functions^9^. CMA-ES generates samples from a multivariate normal distribution where the samples are then ranked using the objective function to be minimized. Based on this ranking, a subset of the samples is then selected and used to update the mean and covariance matrix of the multivariate normal distribution for the next iteration. The process is repeated until convergence. In principle, CMA-ES is equivalent to computing principal component analysis on previously selected samples to direct future sampling. See Hansen^10^ for a full description.

CMA-ES has a number of advantages over alternative methods. As mentioned above, due to its stochastic nature, the algorithm is robust to rough gradients. This is because CMA-ES collects many samples from the multivariate normal distribution that are distributed broadly across the energy landscape rather than centered in one local energy minimum. CMA-ES also takes advantage of updating the covariance matrix which increases the probability of optimal search directions in the next iteration.

The Rosetta macromolecular modeling toolkit is a computational package that offers users and developers state-of-the-art methods for important applications such as structure prediction, protein docking, loop modeling, protein design, antibody modeling and many others^4,11^. Evolution strategies including CMA-ES have recently been applied to structural modeling problems with promising results^12–18^.

Here we implement the CMA-ES algorithm into ROSETTA and evaluate its performance on the problem of peptide docking with constraints. The problem of peptide docking with constraints is important in relation to mimicry of hotspot residues^19–22^. Further, the problem’s simplicity provides an excellent testbed for minimization algorithms and allows the ability to discern performance gains with a manageable number of degrees of freedom. Using the FlexPepDock benchmark of 26 peptides, we compare CMA-ES’s results to Rosetta’s default minimizer as well as a Monte Carlo protocol of small backbone perturbations. We show CMA-ES’s performance is superior using several metrics of structural comparison and show a time analysis demonstrating CMA-ES advantage over the Monte Carlo algorithm.

## Results

### CMA-ES Algorithm Evaluation On Model Tri-peptide

We first implement the CMA-ES algorithm within the ROSETTA framework. We demonstrate the algorithm’s utility by minimizing a model tri-peptide (i.e. Ala-His-Ala) from a high energy state to a lower energy state. The CMA-ES algorithm generates samples from a multivariate distribution. In the case of peptide minimization, the algorithm samples dihedral angles of the peptide backbone and side chains to generate a new conformation (i.e., pose). Once new conformations are generated, each resulting pose is scored using the Rosetta energy function. The poses are ranked according to their score. Top scoring conformations are used to adapt the sampling distribution for the generation of new poses. Supplemental Figure 1A shows conformations from selected iterations throughout the algorithm’s procedure when applied to the tri-peptide. We observe a substantial amount of conformational sampling by iteration 30 allowing the algorithm to explore a rough energy landscape and escape local energy minima. By iteration 60 and iteration 90 sampling focuses on exploring a more local area of the conformational space. And finally, by iteration 120 the algorithm converges toward sampling around the (potentially) global minimum conformation.

We can also observe how individual dihedral angles are being sampled throughout the optimization process. Unlike a Monte Carlo simulation, for example, which is randomly sampling from a static distribution, the CMA-ES algorithm is updating the multivariate distribution every iteration. Therefore, individual dihedral angles may be sampled more or less during optimization and in a correlated manner. We can observe this behavior in Supplemental Figure 1B. We see dihedral angles along the backbone and Histidine side chain of the tri-peptide dynamically change their values as a function of CMA-ES iteration. In particular, we see the first Alanine residue’s Psi angle being sampled continuously through iteration 90 while the Chi angles of the Histidine side chain are relatively constant throughout optimization. This shows that the CMA-ES algorithm efficiently samples only the degrees of freedom necessary to find a low energy conformation.

A major feature of the CMA-ES algorithm is its ability to navigate rough energy landscapes. Consistent with our observations above where we see a large amount of conformational sampling (Supplemental Figure 1A) we also see increases in Rosetta energy at various iterations along the optimization procedure (Supplemental Figure 1C). This increase in conformational sampling and energy allows for the escape out of local energy minima. In particular, we observe a spike in Rosetta energy at iteration 30 where the largest amount of conformational sampling is seen. Rosetta energy is further seen to decrease with less variance until convergence.

### CMA-ES Comparison Evaluation Workflow

In order to compare the performance of the CMA-ES algorithm to other optimization algorithms, we implemented each algorithm into a peptide docking with constraints protocol. Figure 1 shows the overall workflow of the docking protocol and evaluation scheme. As shown in Figure 1A, the first step of the workflow begins with a target peptide bound to a protein. The peptide is extracted from the target keeping the peptide’s conformation fixed (Figure 1B). Next, backbone atoms of the peptide are removed from the pose, leaving only the sidechain atoms (i.e. Disembodied Sidechains, Figure 1C). We then select one sidechain residue arbitrarily to be the primary hotspot and all other side chain residues as ancillary hotspots.

**Figure 1:**
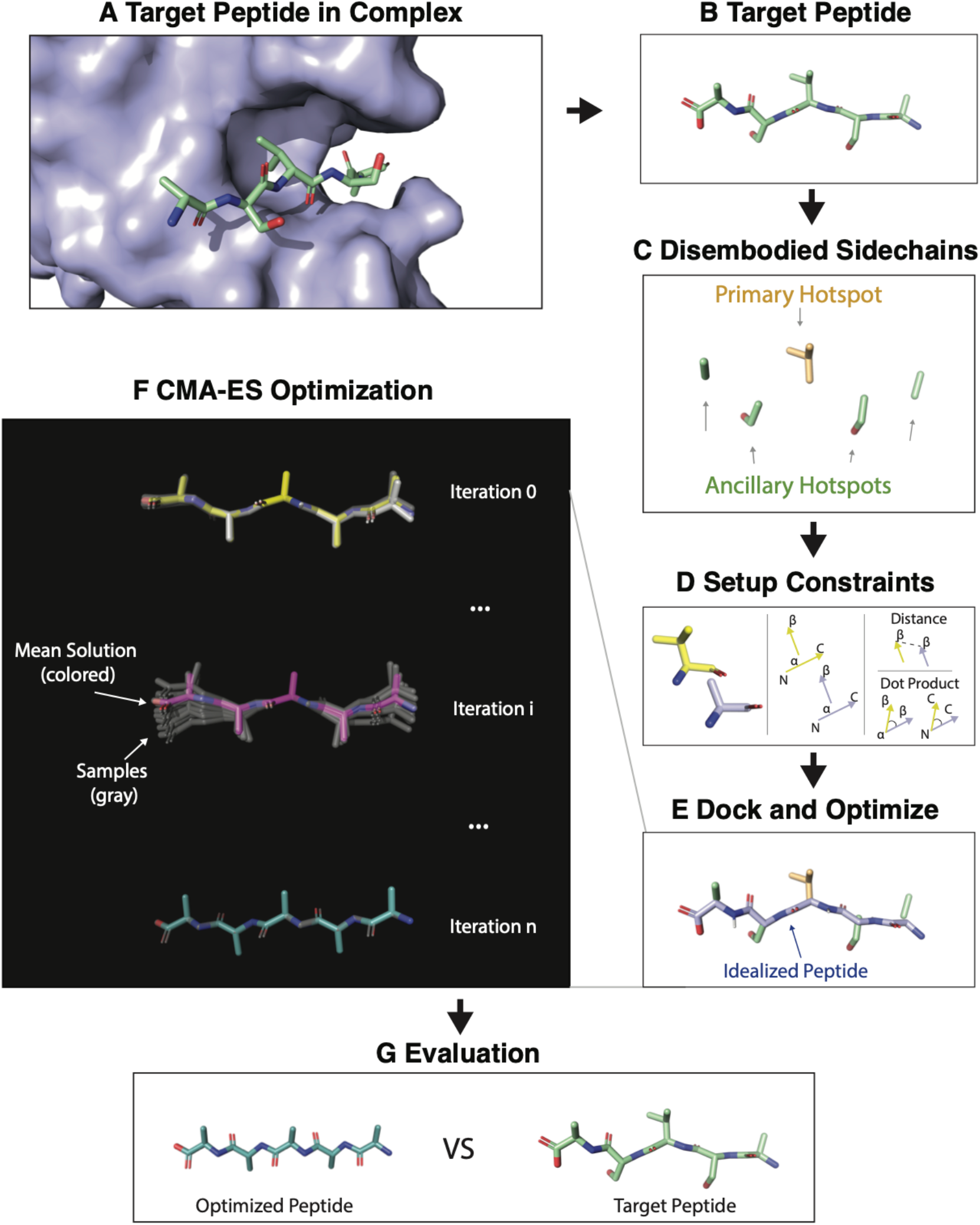
CMA-ES workflow overview. The figure displays a general overview of the CMA-ES test workflow which begins with a peptide (sticks) docked into a protein (surface blue-gray) (A). The target peptide is first isolated from the protein (B). Next, backbone atoms are removed from the target peptide leaving only side chain atoms called disembodied side chains (C). One side chain is selected as the primary hotspot and all others are labeled as ancillary hotspots. Next, constraints are added between residues on an idealized scaffold and the hotspot residues (D). The idealized scaffold is then aligned onto the primary hotspot and phi and psi angles are optimized to match the idealized scaffold residue positions with the ancillary hotspots minimizing the constraints (E). The optimization is done using the CMA-ES optimization protocol where CMA-ES samples each degree of freedom updating the mean and standard deviation of the multivariate normal distribution every iteration (F). Structures of peptides with sampled phi and psi angles are represented in transparent gray. The structure of the updated mean peptide for each iteration is colored. After CMA-ES completes, the final structure is compared to the target peptide for evaluation (G).

Once the hotspots are chosen, an all-alanine idealized peptide scaffold is constructed of the same length and secondary structure of the target peptide. The protocol next puts energy constraints between the atoms of the disembodied side chain residues and the corresponding residues on the idealized scaffold. Constraints are used as described in Fleishman et al.^19^. Each constraint is calculated as a measure of the overlap between the Cβs, the Cα-Cβ vectors, and the C-N vectors of the disembodied residue and the idealized scaffold residue (Figure 1D). The constraint is added to the Rosetta energy function and if fully satisfied will result in a minimum −3 Rosetta Energy Units per residue. The idealized peptide scaffold is then positioned to align to the primary hotspot residue.

The remaining task is to identify phi and psi angle values that align the backbone to the disembodied residues and satisfy the energy constraints. To accomplish this, we implemented the CMA-ES optimization algorithm into the Rosetta framework (Figure 1F). We hypothesized CMA-ES’s ability to traverse rugged energy landscapes would show superior performance on this task compared to gradient based minimization algorithms which get trapped in local minima.

### Comparison of Algorithms on Individual Examples

We applied this workflow to a set of 26 peptides in complex with proteins from the FlexPepDock benchmark^23^. Figure 2 highlights individual examples of peptides. To compare the three algorithms’ performance, we calculated RMSD to the native peptide and Rosetta energy score for the final pose of each individual trajectory and plotted a Rosetta Energy versus RMSD scatterplot. We can see from the example 1NVR in Figure 2A that CMA-ES (blue triangle) outperforms Minimizer (green circle), and Monte Carlo (orange square) sampling algorithms, having lower RMSD to native as well as lower Rosetta Energy. More specifically, we see the top CMA-ES model outperform the Minimizer model by >10 Rosetta Energy Units (REU) and >0.3 RMSD. In Figure 2B, we observe good alignment of the top CMA-ES pose with the native peptide. Further, in Figure 2C, we see 4 out of the 5 hotspot residues constraints are matched with the CMA-ES algorithm. In comparison only 2 are matched with Monte Carlo and 1 (the primary hotspot) is matched using the Minimizer (see Supplemental Table1).

**Figure 2:**
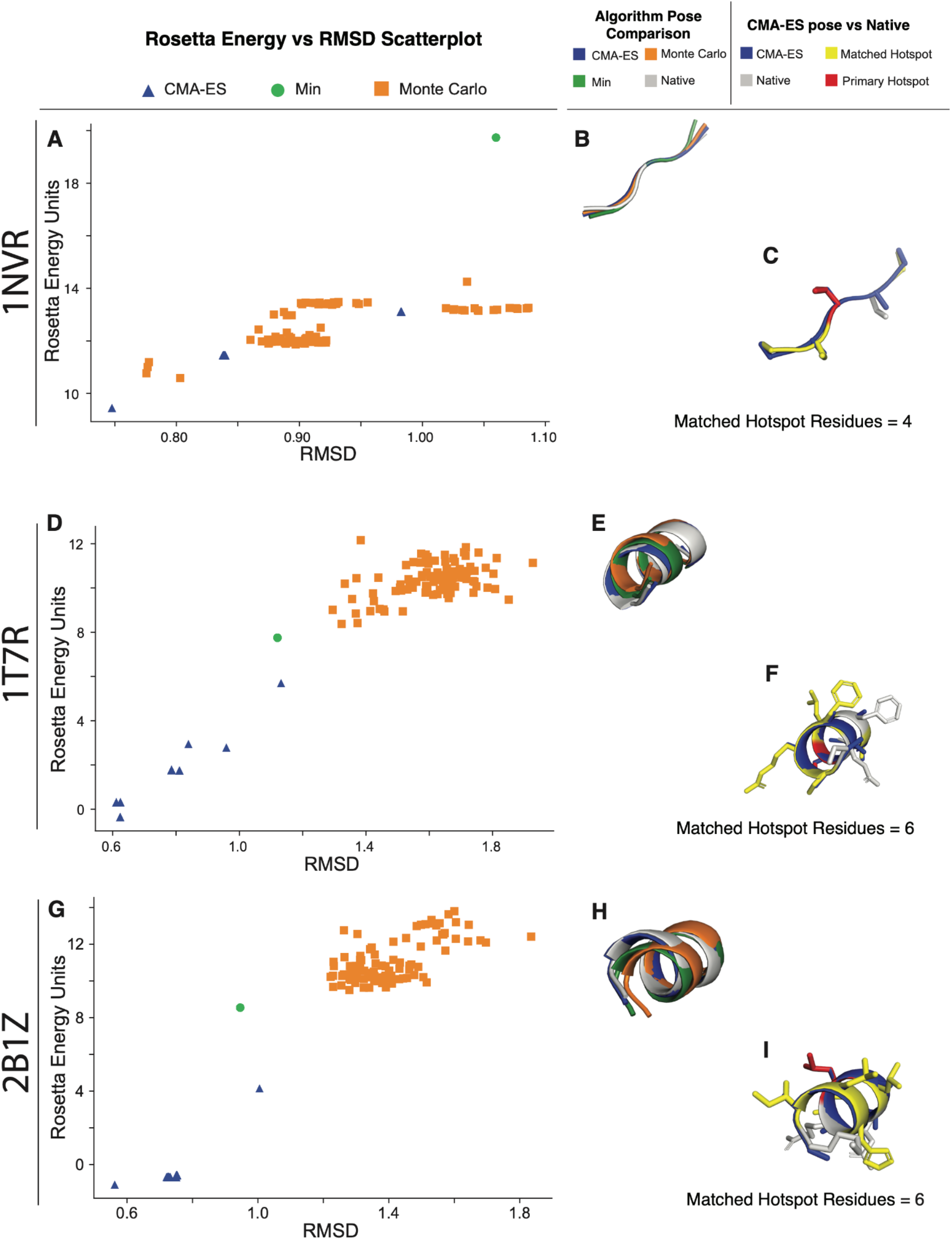
Highlighted examples show CMA-ES outperforms alternative algorithms. Benchmark examples where CMA-ES outperforms Monte Carlo and Gradient-based Minimizer (Min). Row 1: 5-mer peptide from PDB 1NVR. (A) RMSD vs Rosetta Energy scatter plot shows top CMA-ES poses (blue triangles) have lower energy and lower RMSD than poses generated with Monte Carlo (orange squares) or Minimization (green circle). (B) Alignment of lowest energy pose backbones CMA-ES (blue), Monte Carlo (orange), Minimization (green) aligned to native (gray). (C) Alignment of CMA-ES pose to native backbone shows 4 satisfied hotspot constraints (yellow). Primary hotspot is highlighted in red. Row 2: 10-mer peptide from PDB 1T7R. Same colors as row 1. (D) RMSD vs Rosetta Energy scatter plot shows CMA-ES poses have lower energy and lower RMSD than Monte Carlo and Minimization. (E) Lowest energy CMA-ES pose aligns very well to native compared to other lowest energy alternative algorithm poses. (F) CMA-ES pose aligned to native satisfies 6 hotspot constraints. Row 3:9-mer peptide from PDB 2B1Z. Same colors as row 1. (G) RMSD vs Rosetta Energy scatter plot shows nearly all CMA-ES poses have lower energy and lower RMSD than Monte Carlo and Minimization. (H) Lowest energy CMA-ES pose aligns well with native backbone compared to other lowest energy alternative algorithm poses. (I) CMA-ES pose aligned to native satisfies 6 hotspot constraints.

Similar to 1NVR, we see CMA-ES outperform the Minimizer and Monte Carlo algorithms on other examples as well. In Figure 2D and 2G CMA-ES shows superior performance versus the other algorithms, improving on both energy (REU) and RMSD for peptides in PDBs 1T7R and 2B1Z respectively. When we inspect the poses overlaid with the native peptides (Figure 2E and 2H), we see a substantial improvement in terms of RMSD. The CMA-ES poses (blue) match very well to the native (gray) while the Minimizer (green) and Monte Carlo (orange) produce poses that deviate from the native. We also identify 6 matched hotspot residues by CMA-ES for each example, 1T7R and 2B1Z (Figure 2F and 2I). This is substantially better than the Minimizer and Monte Carlo algorithms which had at most 3 matched hotspot residues but often lower (see Supplemental Table1). Additional benchmark examples can be found in Supplemental Figure 2 which show CMA-ES improved performance over the other algorithms.

### Global Comparison of Algorithms

Now that we have established evidence of individual examples of CMA-ES outperforming other optimization algorithms, we evaluate each algorithm’s models on all 26 peptides in the benchmark for a global comparison. Figure 3A shows a bar graph of the number of benchmark instances where each algorithm matched more hotspot residues than the other algorithms. In this winner-take-all analysis, CMA-ES matched more hotspot residues than the other algorithms for 11 benchmark instances while Monte Carlo and Minimizer produced the top model for only 2 and 1 benchmark instances, respectively. Alternatively, we evaluate the difference of the number matched hotspot residues between all three algorithms. This analysis provides a quantitative look at not just which algorithm had the top scoring model but by how much. In Supplemental Figure 3A, we observe many benchmark instances where the CMA-ES algorithm matched +2, +3, and +4 hotspot residues relative to the Monte Carlo algorithm while there were no instances where the Monte Carlo algorithm matched greater than +2 compared to CMA-ES. Similar behavior can be observed in Supplemental Figure 3B, where many instances of CMA-ES match > +2 compared to the Minimizer algorithm while no instances were observed where the Minimizer matched more than +2 in comparison to CMA-ES. We also evaluate the Monte Carlo algorithm versus the Minimizer algorithm (Supplemental Figure 3C) and observe the Monte Carlo algorithm performs better.

**Figure 3:**
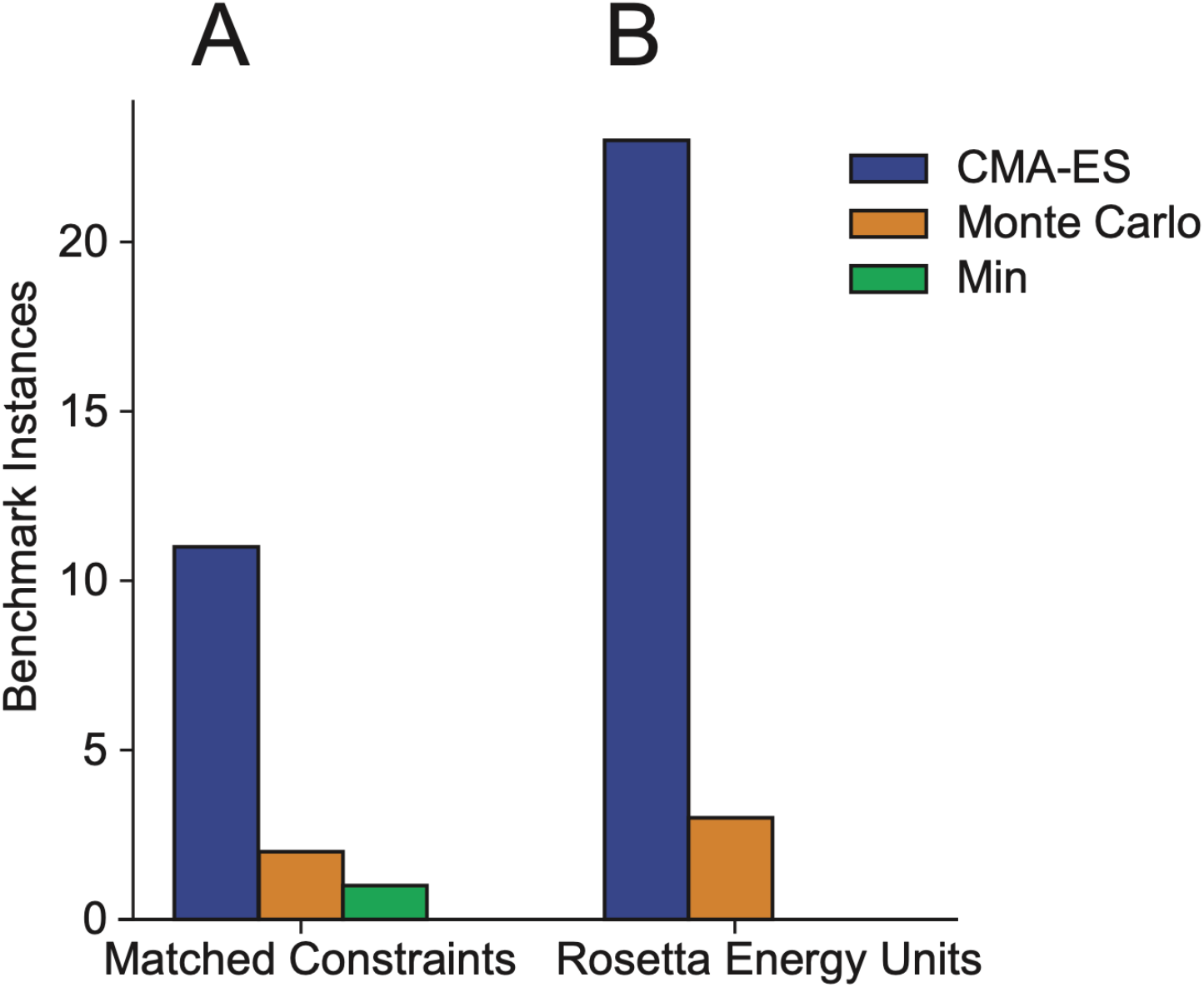
Global benchmark analysis showing CMA-ES performance. The bar plot shows the number of benchmark instances where CMA-ES (blue), Monte Carlo (orange), or Minimizer (green) algorithms produced the top performing model. (A) Evaluation of top performing models based on the number of matched hotspot residues. CMA-ES matched more hotspot residues than the other algorithms for 11 benchmark instances while Monte Carlo and Minimizer produced the top model for only 2 and 1 benchmark instances, respectively. (B) Evaluation of top performing models based on Rosetta Energy. CMA-ES models scored lower energies for 23 benchmark instances compared to only 3 for Monte Carlo and 0 for Minimizer.

In addition to matching hotspot residues, we also evaluate each algorithms’ performance in regards to total Rosetta energy. Figure 3B shows the CMA-ES models scored lower energies for 23 benchmark instances compared to only 3 for Monte Carlo and 0 for Minimizer. Supplemental Figure 3D shows the difference in Rosetta energy between models produced by CMA-ES and Monte Carlo algorithms. The mean difference between Rosetta energies of CMA-ES and Monte Carlo algorithms is −2.1 REU but as the plot shows, for some benchmark instances, the CMA-ES models scored substantially lower in energy compared to Monte Carlo models (~10 REU). When we compare CMA-ES to the Minimizer algorithm, we see a more pronounced effect (Supplemental Figure 3E) with CMA-ES models having substantially lower Rosetta energies including a mean difference in Rosetta energies of −12.6. Finally, in Supplemental Figure 3F, we observe the Monte Carlo algorithm having substantially lower Rosetta energies when compared directly to the Minimizer algorithm.

We finally evaluate the algorithms in terms of their models’ root mean square distance (RMSD) to the native peptide. We observe similar performance among the algorithms. In particular, we see the algorithms perform within +/- 0.5 RMSD of each other on the majority of benchmark instances (Supplemental Figure 3G-I). We do note however that the Monte Carlo algorithm shows a slightly better performance than CMA-ES (Supplemental Figure 3G). This is likely due to the Monte Carlo algorithm’s sampling from a Ramachandran map probability distribution based on experimental solved structures. This would allow the Monte Carlo algorithm to recapitulate native conformations. A direct comparison of CMA-ES with the Minimizer algorithm, however, shows the majority of CMA-ES models are more similar to the native peptide (Supplemental Figure 3H). Lastly, a direct comparison of the Monte Carlo and Minimizer algorithm shows the majority of Monte Carlo models are closer to the native peptide in terms of RMSD (Supplemental Figure 3I).

Using these standard performance metrics, we see CMA-ES is a robust optimization algorithm. We see that it outcompetes both the standard default Rosetta Minimizer and a customized Monte Carlo algorithm using Rosetta energy score. It also outcompetes the Minimizer in terms of RMSD to the native peptide. Finally, CMA-ES matches far more hotspot residues than either of the other algorithms.

### Time Comparison of Algorithms

With most optimization algorithms, there is a tradeoff between speed and sampling. We next evaluate the algorithms in terms of their run time to evaluate their efficiency to better understand this tradeoff. As can be seen in Figure 4, the Rosetta Minimizer has the fastest runtime with a median runtime of 15 seconds among all benchmark instances. The CMA-ES algorithm has the second fastest runtime with a median of 59 seconds while the Monte Carlo algorithm has an order of magnitude longer median time of 695 seconds. We therefore conclude based on the time analysis that the CMA-ES algorithm efficiently samples the energy landscape and effectively identifies low energy conformations appropriately balancing the speed/sampling tradeoff.

**Figure 4:**
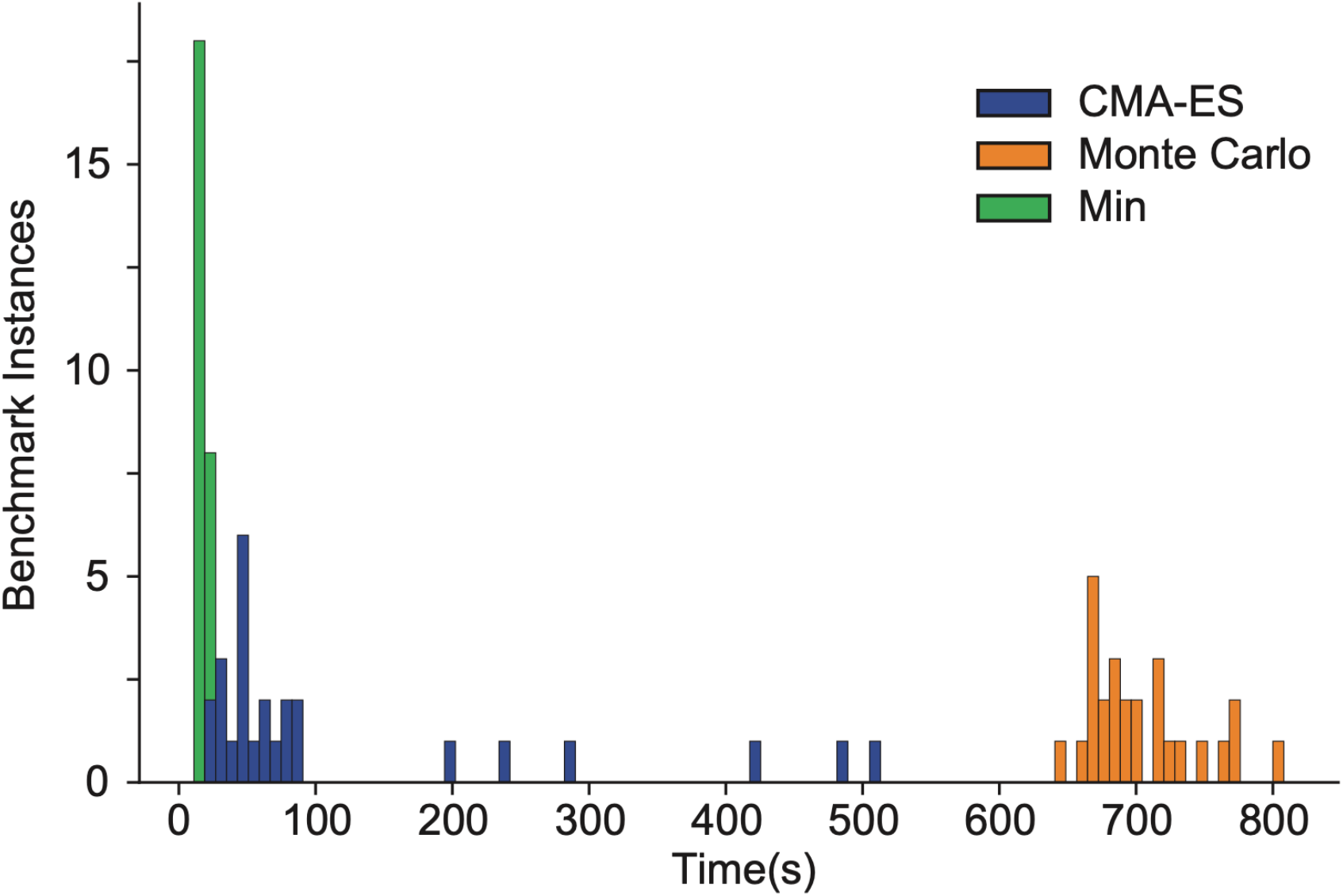
Run Time Analysis demonstrates CMA-ES is competitive with Minimization algorithm. Histograms of runtimes for benchmark instances run using each optimization algorithm: CMA-ES (blue), Monte Carlo (orange), and Minimization (green). Minimization had a median runtime of 15 seconds, while CMA-ES had a median runtime of 59 seconds. Monte Carlo had a substantially slower median runtime of 695 seconds.

## Methods

### CMA-ES implementation in Rosetta

We based our Rosetta CMA-ES optimization implementation on the core c-cmaes library code from^10^ (https://github.com/CMA-ES/c-cmaes). The CMA-ES optimization algorithm is first initialized with a vector of degrees of freedom which takes the form of initial means of the multivariate normal distribution. Similar to other minimization algorithms implemented in Rosetta, degrees of freedom represent torsion angles along a peptide backbone (i.e. phi, psi, and omega), side chain torsion angles (ie. chi), and rotational/translational degrees of freedom. A vector of sigma values is also initialized representing the starting standard deviation of each degree of freedom in the multivariate normal distribution. We use the default value of 0.3 for the starting standard deviation.

Once initialized, the algorithm then obtains a population of samples from the multivariate normal distribution of size lambda. We use the recommended lambda value of 4+int(3*log(N)), where N is the number of degrees of freedom. Each sample is evaluated using the selected Rosetta energy score function (ref2015 for this application). The samples are then ranked based on their evaluated energies and the top mu samples are then used to select the new mean and update the covariance matrix of the next iteration’s multivariate normal distribution. Mu is set to ½ of lambda. The algorithm iterates until either a selected tolerance parameter of 0.001 is reached or a max number of iterations of 1000 is reached.

All associated code is freely accessible to academic users via the ROSETTACOMMONS website (http://www.rosettacommons.org).

### Monte Carlo Algorithm

We used the Monte Carlo algorithm implementation in Rosetta which is a modification of the protocol used in Renfrew et al. 2011^24^. Briefly, the protocol iterates through a perturbation cycle ten times, randomly making small and shear moves of backbone phi and psi angles of max 2 degrees. All phi and psi angles along the peptide backbone are subject to perturbation. Monte Carlo temperature for perturbation moves is set at 0.8 while each iteration temperature value is set at 0.5. In contrast to the original algorithm in Renfrew et al. 2011, docking (i.e. rotation and translation) is turned off in order to keep the scaffold fixed on the primary hotspot residue. Also, the design step is turned off as well as no final design minimization step. We generated 100 models.

### Minimization Algorithm

We used the default gradient-based minimizer in Rosetta which optimizes a set of degrees of freedom for a specified score function. The default minimizer is an implementation of the L-BFGS algorithm (i.e. lbfgs_armijo_nonmonotone)^25^. We allowed only backbone torsion angles to move. We set the tolerance parameter to 0.001.

### Model Tri-peptide CMA-ES Minimization

We used pymol to construct an Ala-His-Ala peptide. We used the commandline below to minimize the model tri-peptide.

`./Rosetta/main/source/bin/minimize.default.linuxgccrelease-database`

./Rosetta/main/database/-s./AHA.pdb-run:min_type **cmaes**

### Benchmark Workflow

We downloaded structures of peptide-protein interaction pairs found in the FlexPepDock benchmark (https://doi.org/10.1371/iournal.pone.0018934.s002)^23^. Each structure was “cleaned” using the pdb_clean.py script provided in the Rosetta tools directory. Structures were then “relaxed” using Rosetta’s FastRelax protocol with the ‘-relax:constrain_relax_to_start_coords’ flag. 50 decoys were produced and the lowest scoring model was selected. For each benchmark instance, the peptide was extracted from the target and backbone atoms removed, resulting in a set of disembodied sidechains. Glycine residues were ignored. Each disembodied sidechain was placed in a separate pdb formatted stub constraint library file. Idealized peptides of the same size and nearest secondary structure as the target peptide were generated using PyMOL^26^. A primary hotspot residue was selected for each peptide and all other disembodied side chains are considered ancillary hotspot residues. The idealized backbone is aligned to the primary hotspot residue. Backbone stub constraints are created for the primary hotspot and all ancillary hotspot residues. 100 decoy samples were run for each peptide and each algorithm (i.e. Monte Carlo, Minimization, and CMA-ES). The Rosetta score function ref2015 was used for all energy calculations.

## Discussion

Herein, we describe the implementation of a blackbox optimization algorithm (CMA-ES) into the Rosetta macromolecular modeling framework. We first demonstrate the algorithm’s utility by minimizing the energy of a model tri-peptide. We show CMA-ES is capable of escaping local energy minima while lowering the overall energy. In addition, we observe CMA-ES efficiently samples degrees of freedom so as to not waste computation on unnecessary sampling.

We also analyze CMA-ES performance on the task of docking peptide scaffolds to hotspot residues from a benchmark of 26 peptides. We demonstrate CMA-ES outperforms gradientbased minimization and Monte Carlo algorithms on several evaluation metrics including total number of matched hotspot residues and overall decrease in energy. Interestingly, we observed the Monte Carlo algorithm slightly outperform CMA-ES with respect to RMSD to native (Supplemental Figure 3G). This result may be due to the Monte Carlo sampling procedure using Rosetta small and shear moves which sample from the Ramachandran distribution of phi and psi angles. Sampling from the Ramachandran distribution is likely to ensure Monte Carlo poses are in a more peptide-like conformation. CMA-ES on the other hand samples from a multivariate normal distribution across all degrees of freedom and therefore has no prior information on peptide geometry other than when a pose is evaluated by the Rosetta energy function. Minor inaccuracies in the Rosetta energy function may be the cause, and further investigation is needed to determine the source.

Finally, we evaluated all three algorithms in terms of their runtime. While gradient-based minimization performed the fastest as expected, CMA-ES was highly competitive on the broad majority of the test cases (Figure 4). The Monte Carlo algorithm performed slowest as expected. Energy optimization is often a tradeoff between speed and finding a lower energy minima. CMA-ES balances this tradeoff by consistently finding lower energy minima than both the gradient based minimization algorithm and Monte Carlo algorithm as well as being faster than the Monte Carlo algorithm.

Unlike other evolutionary algorithms, CMA-ES does not require many parameters to be specified at runtime. A user only provides an initial starting conformation and initial sigma value. The sigma value is automatically updated after each iteration to determine optimal values, so the algorithm is robust to initial values. A population value lambda can be modified but the default value which is a function of the number of degrees of freedom has been shown to work well in most cases. Additional strategies are being explored to increase the population size after restart which has shown additional performance improvements^27^. Moreover, hybrid strategies that combine gradient-based optimization and evolution strategies also show great promise in boosting performance^28^. Overall however, the algorithm in its current state requires minimal to no tuning from the user. This points to its high-value in many future modeling applications including ligand docking^2^, peptidomimetic modeling^29^ and design^20^, and cyclic peptide design^30^.

## Acknowledgements

This work was supported by grants from the NIH (R00 HD092613 and L40 HD096554 to K.D.) and RosettaCommons (RC22024 to K.D.). The authors would also like to thank José Villegas and Parisa Hosseinzadeh for helpful discussions.

## Supplemental Figures

**Supplemental Figure 1:** Demonstration of CMA-ES algorithm applied to model Ala-His-Ala tripeptide. (A) Snapshot images of tri-peptide at subsequent iterations throughout the optimization procedure. CMA-ES samples a multivariate distribution based on the degrees of freedom of the tri-peptide. Transparent gray conformations represent samples and yellow conformation represents mean of sample population. Early iterations have increased conformational sampling (iteration 30). Sampling begins to stabilize around iteration 120 and ultimately converges. (B) Heatmap displays the value of dihedral angles during the CMA-ES optimization procedure. Similar to the observation in (A), substantial sampling is seen around iteration 30. The first Alanine’s Psi angle is notably variable past iteration 90. Histidine Chi1, and Chi2 angles are mostly stable throughout the procedure. Omega angles which change minimally throughout the run are omitted for clarity. (C) Graph displays Rosetta energy units at each iteration of the CMA-ES run. Consistent with (A), Rosetta energy is highly variable at iteration 30 where CMA-ES undergoes the most sampling. The energy values subsequently drop and become substantially less variable after iteration 80.

**Supplemental Figure 2: All Benchmark RMSD vs Rosetta Energy Scatterplots and Pose Alignment to Native.** Each row represents a benchmark peptide. Left column: Scatterplots of RMSD vs Rosetta Energy Units showing evaluation of 100 poses each produced by CMA-ES (blue triangles), Monte Carlo (orange squares), and Minimization (green circle). Right column: Alignment of lowest energy pose backbones CMA-ES (blue), Monte Carlo (orange), Minimization (green) aligned to native (gray).

**Supplemental Figure 3: Pairwise Algorithm Metric Comparison.** Distribution of differences between algorithms for individual evaluation metrics. Each data point represents an individual benchmark instance. All metrics were calculated on lowest energy poses from respective algorithms. Box plots with quartile ranges are plotted inside each kernel density violin plot. (A-C) Difference in number of matched hotspot residues. (A) CMA-ES vs Monte Carlo shows CMA-ES (blue) matched more hotspot residues than Monte Carlo (orange). Equal number matched (black). (B) CMA-ES vs Minimization shows CMA-ES matched more hotspot residues than Minimization (green). (C) Minimization vs Monte Carlo shows Monte Carlo matched more hotspot residues than Minimization. (D-F) Difference in Rosetta Energy. (D) CMA-ES vs Monte Carlo shows CMA-ES has lower energy than Monte Carlo. (E) CMA-ES vs Minimization shows CMA-ES has lower energy than Minimization. (F) Minimization vs Monte Carlo shows Monte Carlo has lower energy than Minimization. (G-I) Difference in root mean squared deviation (RMSD) to the native peptide. (G) CMA-ES vs Monte Carlo shows the majority of Monte Carlo poses are closer to native than CMA-ES. (H) CMA-ES vs Minimization shows most CMA-ES poses are closer to native than Minimization. (I) Minimization vs Monte Carlo shows most Monte Carlo poses are closer to native than Minimization.

## Supplemental Tables

**Supplemental Table 1: Benchmark evaluation metric table.**

## Supplemental Files

**Supplemental File 1 (zip): PDB formatted benchmark inputs and top models.**

## References

1. Jumper, J. et al. Highly accurate protein structure prediction with AlphaFold. Nature 596, 583–589 (2021).

2. Fu, D. Y. & Meiler, J. RosettaLigandEnsemble: A Small-Molecule Ensemble-Driven Docking Approach. ACS Omega 3, 3655–3664 (2018).

3. Chevalier, A. et al. Massively parallel de novo protein design for targeted therapeutics. Nature 550, 74–79 (2017).

4. Leman, J. K. et al. Macromolecular modeling and design in Rosetta: recent methods and frameworks. Nat. Methods 17, 665–680 (2020).

5. Polyak, B. Introduction to Optimization. (2020).

6. Onuchic, J. N., Luthey-Schulten, Z. & Wolynes, P. G. THEORY OF PROTEIN FOLDING: The Energy Landscape Perspective. Annu. Rev. Phys. Chem. 48, 545–600 (1997).

7. Kuhlman, B. & Bradley, P. Advances in protein structure prediction and design. Nat. Rev. Mol. Cell Biol. 20, 681–697 (2019).

8. Li, Z., Lin, X., Zhang, Q. & Liu, H. Evolution strategies for continuous optimization: A survey of the state-of-the-art. Swarm Evol. Comput. 56, 100694 (2020).

9. Hansen, N. & Ostermeier, A. Completely Derandomized Self-Adaptation in Evolution Strategies. Evol. Comput. 9, 159–195 (2001).

10. Hansen, N. The CMA Evolution Strategy: A Comparing Review. 28.

11. Leaver-Fay, A. et al. Rosetta3: An Object-Oriented Software Suite for the Simulation and Design of Macromolecules. Methods Enzymol. 487, 545–574 (2011).

12. Rakhshani, H., Idoumghar, L., Lepagnot, J. & Brévilliers, M. Speed up differential evolution for computationally expensive protein structure prediction problems. Swarm Evol. Comput. 50, 100493 (2019).

13. Winter, R. et al. Efficient multi-objective molecular optimization in a continuous latent space. Chem. Sci. 10, 8016–8024 (2019).

14. Maximova, T., Moffatt, R., Ma, B., Nussinov, R. & Shehu, A. Principles and Overview of Sampling Methods for Modeling Macromolecular Structure and Dynamics. PLOS Comput. Biol. 12, e1004619 (2016).

15. Desjarlais, J. R. & Handel, T. M. De novo design of the hydrophobic cores of proteins. Protein Sci. Publ. Protein Soc. 4, 2006–2018 (1995).

16. Raha, K., Wollacott, A. M., Italia, M. J. & Desjarlais, J. R. Prediction of amino acid sequence from structure. Protein Sci. Publ. Protein Soc. 9, 1106–1119 (2000).

17. Clausen, R., Sapin, E., De Jong, K. A. & Shehu, A. Evolution Strategies for Exploring Protein Energy Landscapes. in Proceedings of the 2015 Annual Conference on Genetic and Evolutionary Computation 217–224 (Association for Computing Machinery, 2015). doi:10.1145/2739480.2754692.

18. Müller, C. L. & Sbalzarini, I. F. Energy landscapes of atomic clusters as black box optimization benchmarks. Evol. Comput. 20, 543–573 (2012).

19. Fleishman, S. J. et al. Computational design of proteins targeting the conserved stem region of influenza hemagglutinin. Science 332, 816–821 (2011).

20. Lao, B. B. et al. Rational Design of Topographical Helix Mimics as Potent Inhibitors of Protein–Protein Interactions. J. Am. Chem. Soc. 136, 7877–7888 (2014).

21. Drew, K. et al. Adding Diverse Noncanonical Backbones to Rosetta: Enabling Peptidomimetic Design. PLOS ONE 8, e67051 (2013).

22. Thanos, C. D., DeLano, W. L. & Wells, J. A. Hot-spot mimicry of a cytokine receptor by a small molecule. Proc. Natl. Acad. Sci. U. S. A. 103, 15422–15427 (2006).

23. Raveh, B., London, N., Zimmerman, L. & Schueler-Furman, O. Rosetta FlexPepDock ab-initio: Simultaneous Folding, Docking and Refinement of Peptides onto Their Receptors. PLOS ONE 6, e18934 (2011).

24. Renfrew, P. D., Choi, E. J., Bonneau, R. & Kuhlman, B. Incorporation of Noncanonical Amino Acids into Rosetta and Use in Computational Protein-Peptide Interface Design. PLOS ONE 7, e32637 (2012).

25. Liu, D. C. & Nocedal, J. On the limited memory BFGS method for large scale optimization. Math. Program. 45, 503–528 (1989).

26. The PyMOL Molecular Graphics System, Version 2.0 Schrodinger, LLC.

27. Auger, A. & Hansen, N. A restart CMA evolution strategy with increasing population size. in 2005 IEEE Congress on Evolutionary Computation vol. 2 1769–1776 Vol. 2 (2005).

28. Maheswaranathan, N., Metz, L., Tucker, G., Choi, D. & Sohl-Dickstein, J. Guided evolutionary strategies: augmenting random search with surrogate gradients. in Proceedings of the 36th International Conference on Machine Learning 4264–4273 (PMLR, 2019).

29. Butterfoss, G. L. et al. De novo structure prediction and experimental characterization of folded peptoid oligomers. Proc. Natl. Acad. Sci. 109, 14320–14325 (2012).

30. Hosseinzadeh, P. et al. Anchor extension: a structure-guided approach to design cyclic peptides targeting enzyme active sites. Nat. Commun. 12, 3384 (2021).

