## Supplementary figures and images for "CMA-ES-Rosetta: Blackbox optimization algorithm traverses rugged peptide docking energy landscapes"

### Supplemental Figure 1

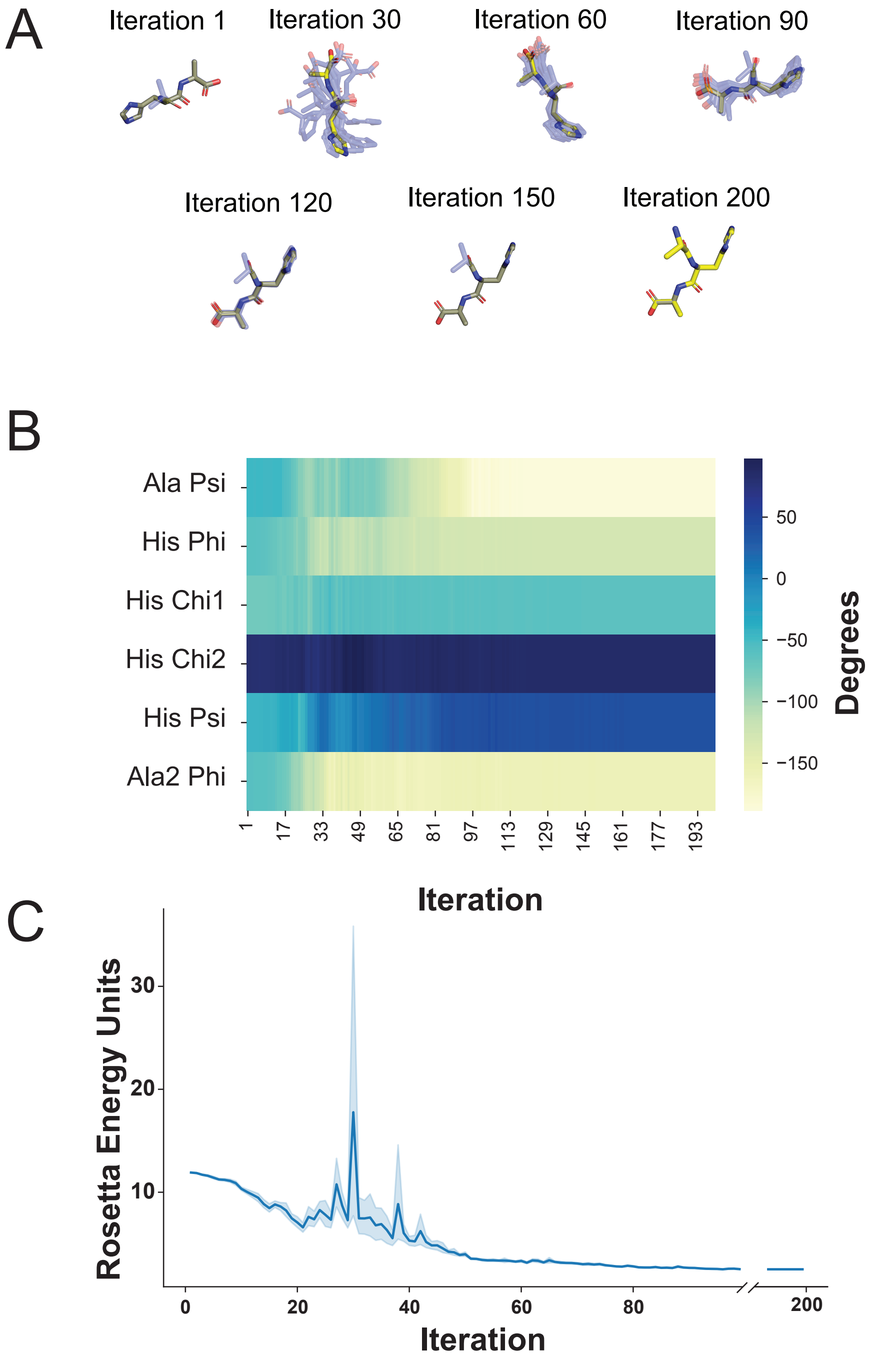

### Supplemental Figure 2

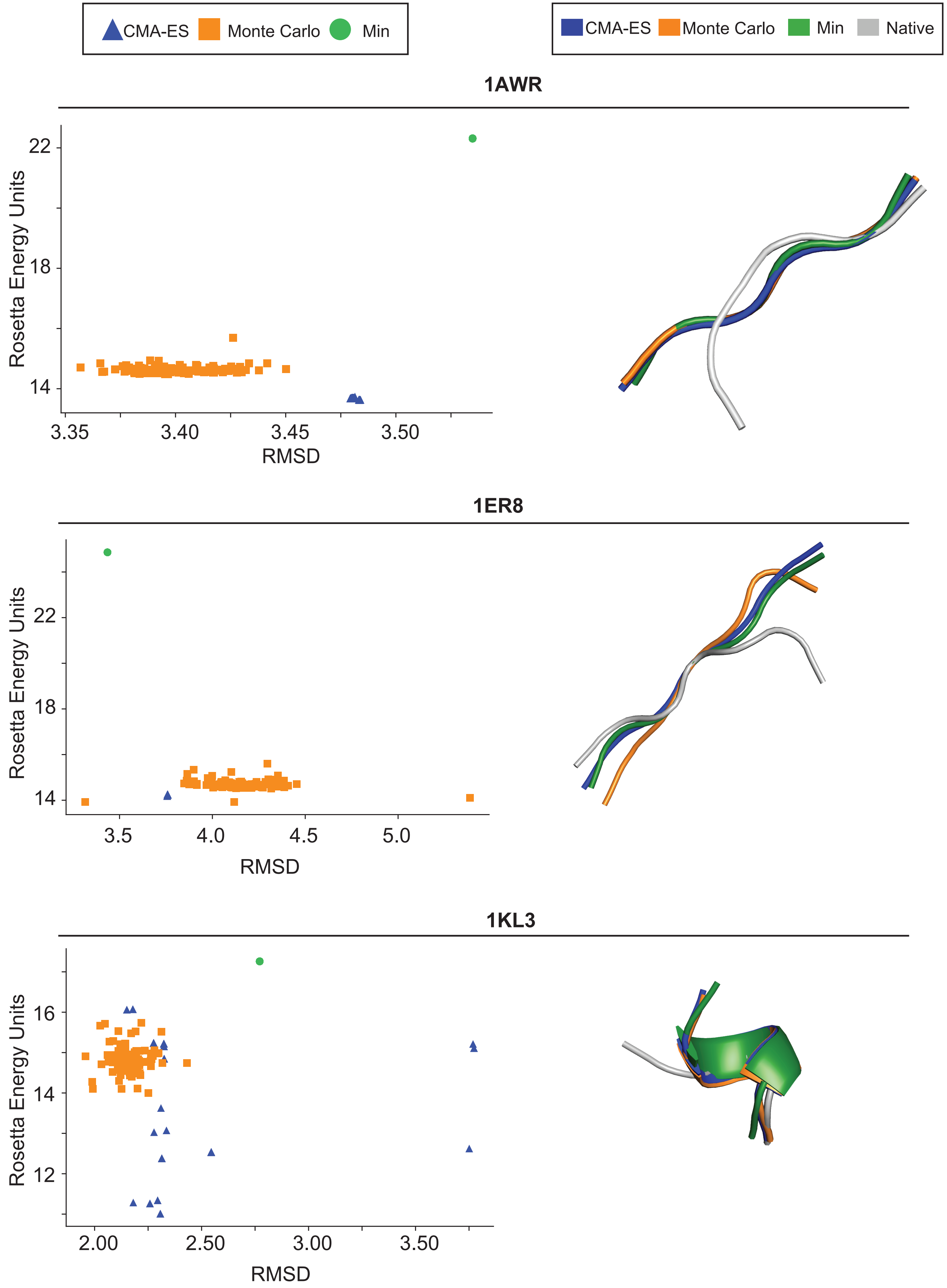

**1N7F**

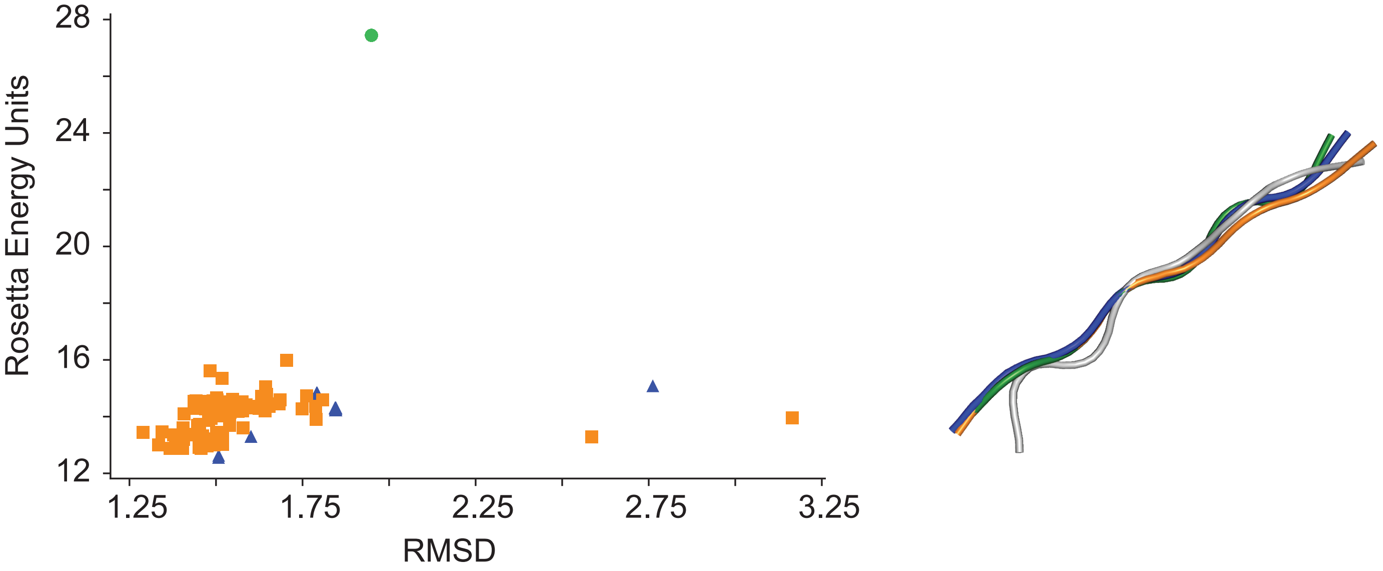


**1NLN**

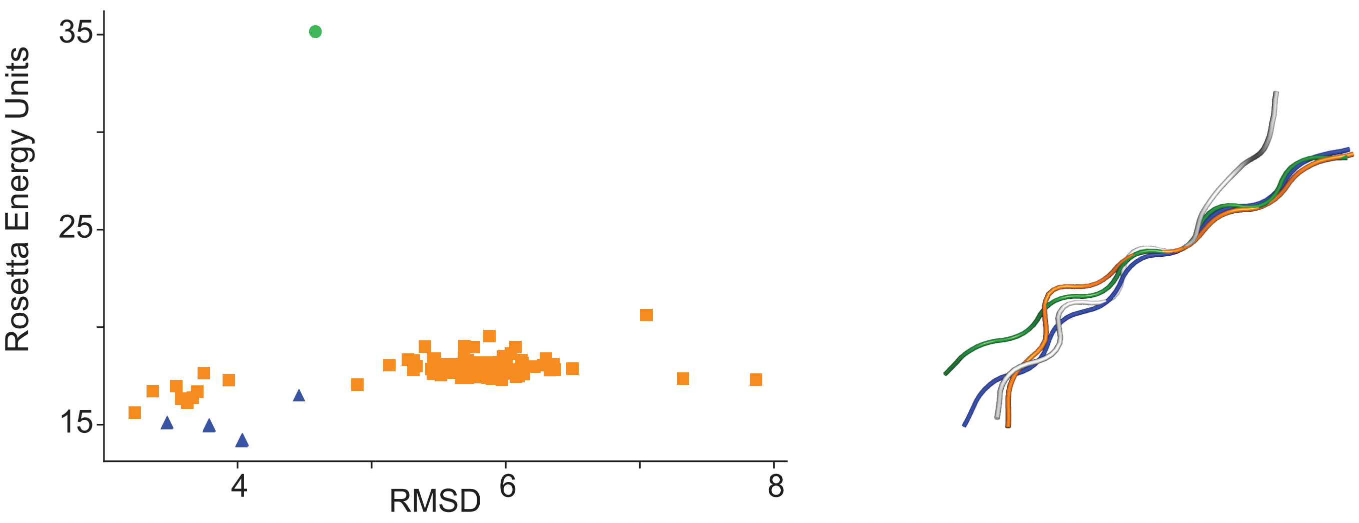


**1QKZ**

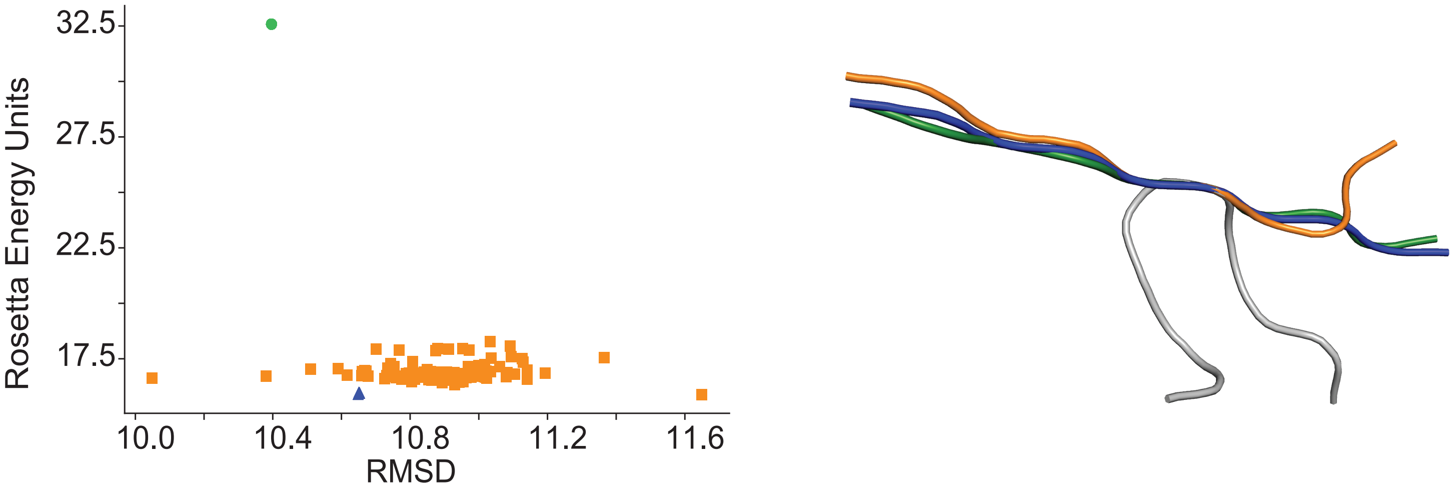

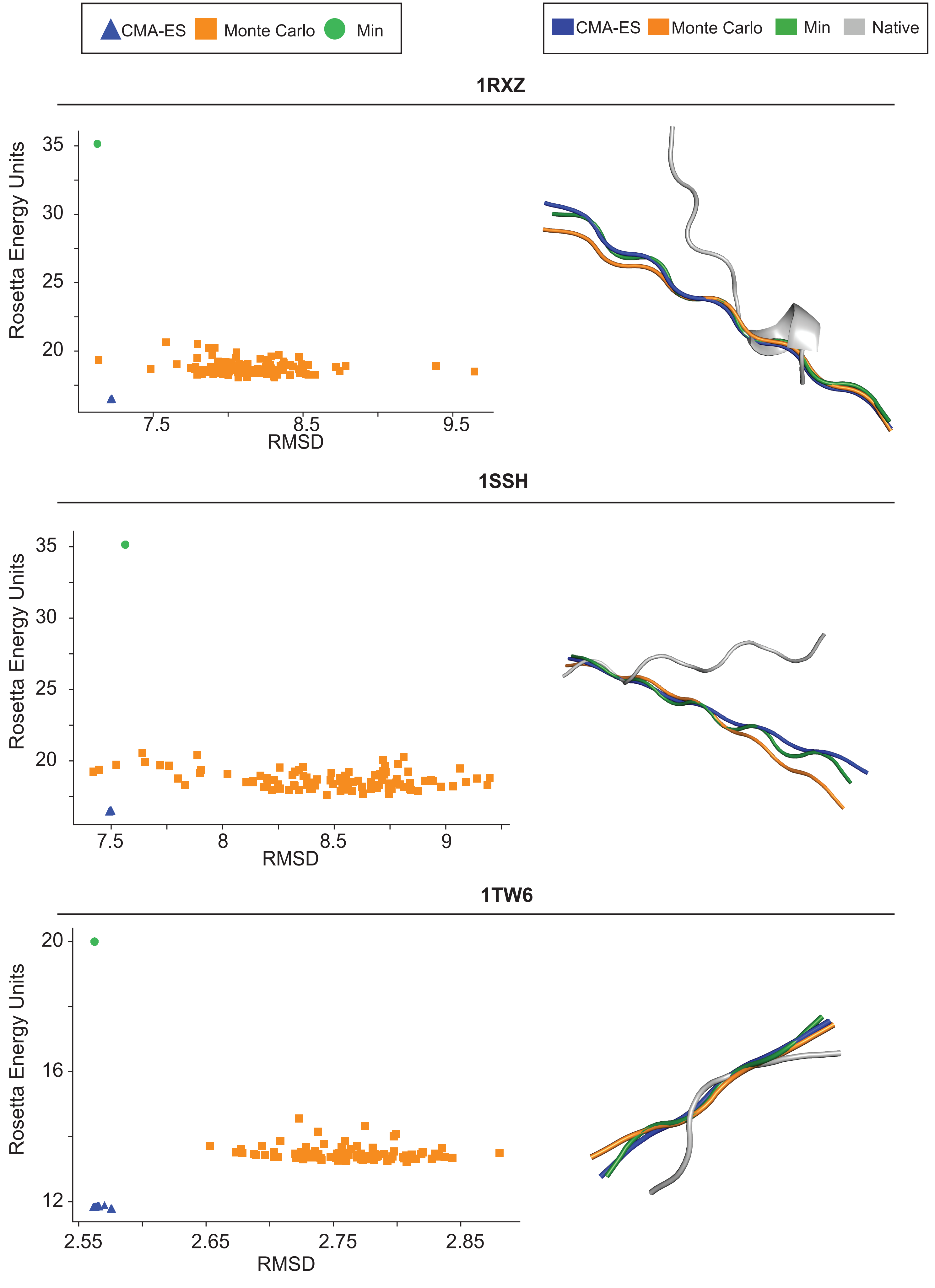

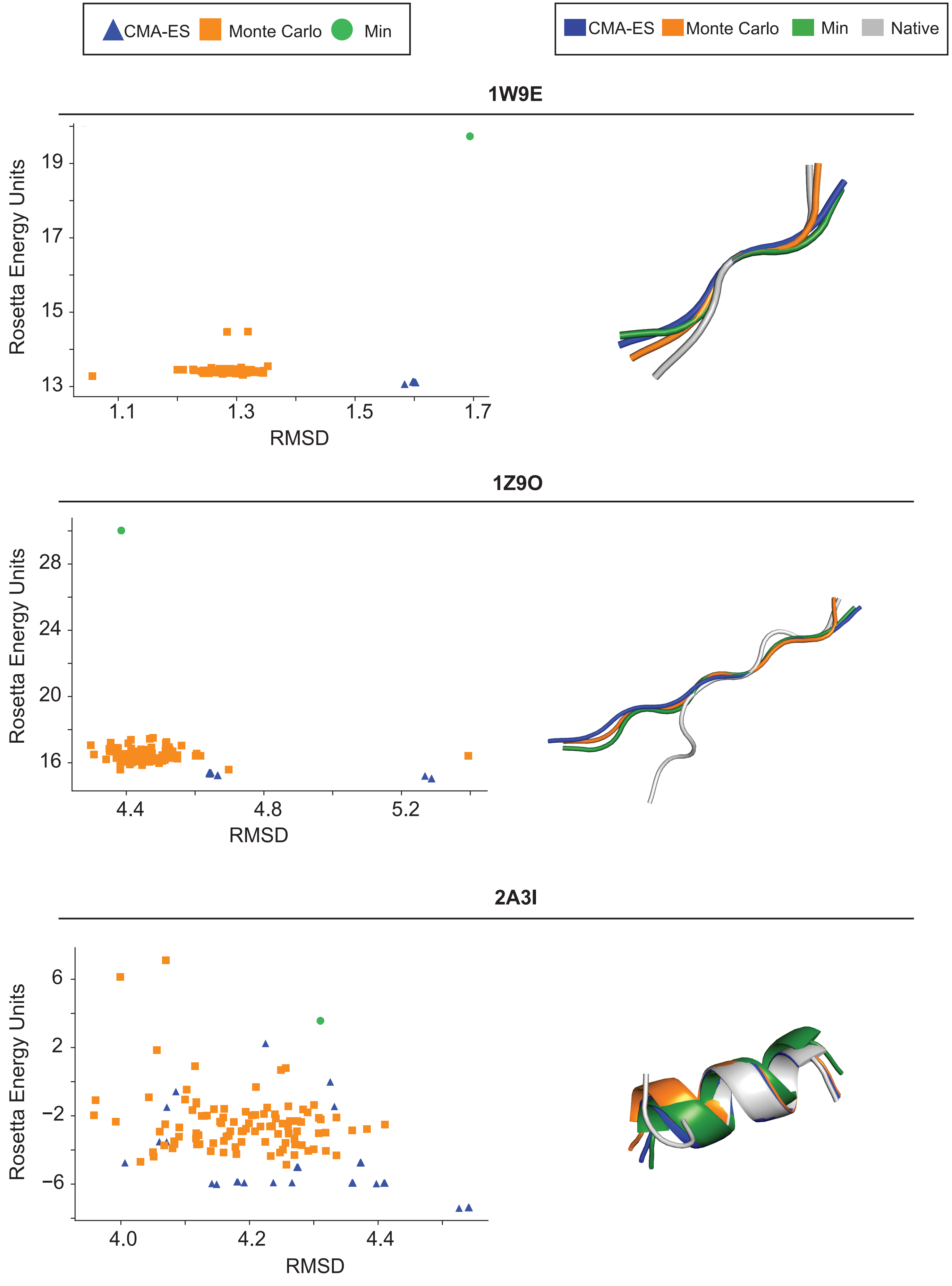

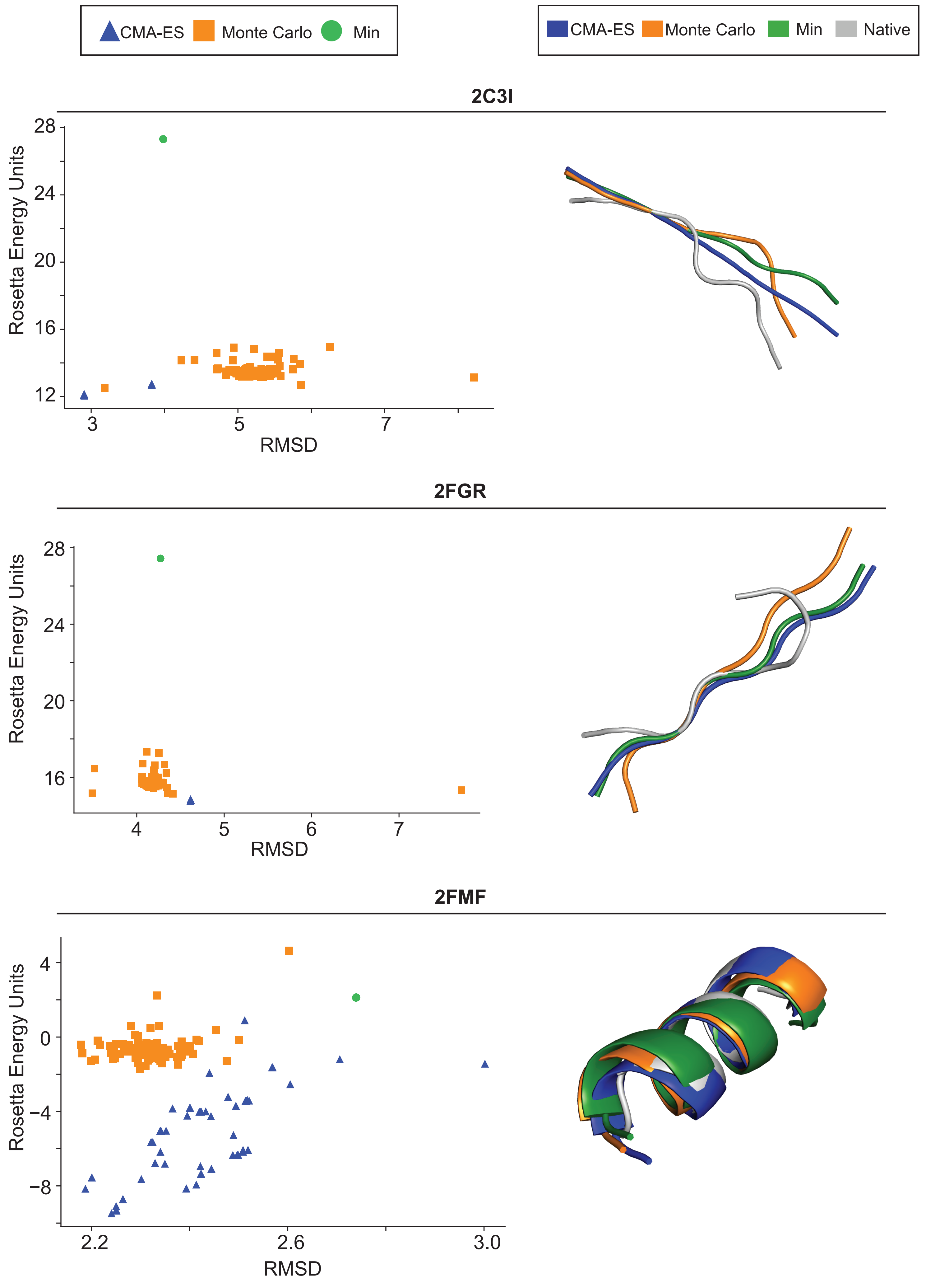

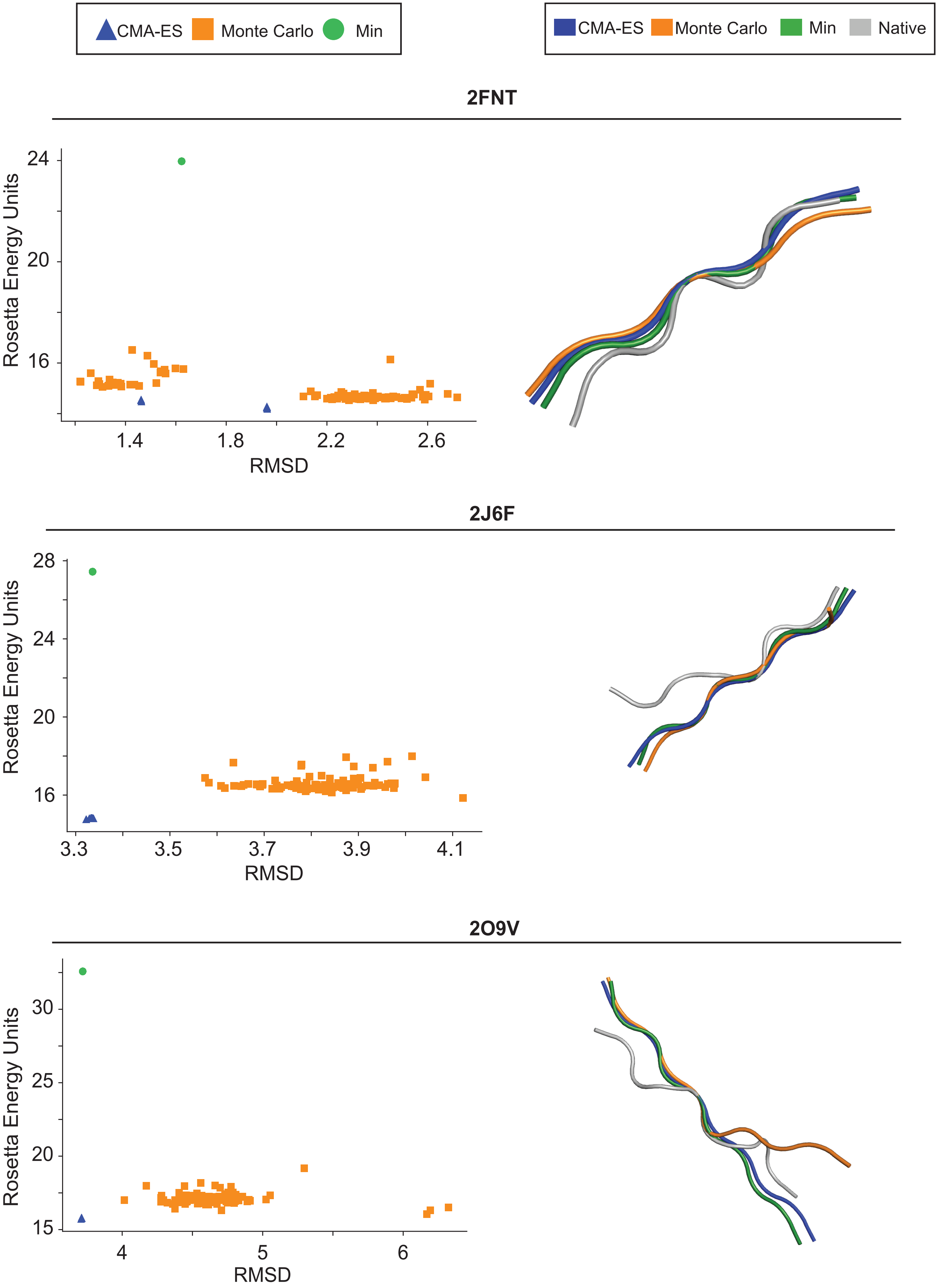

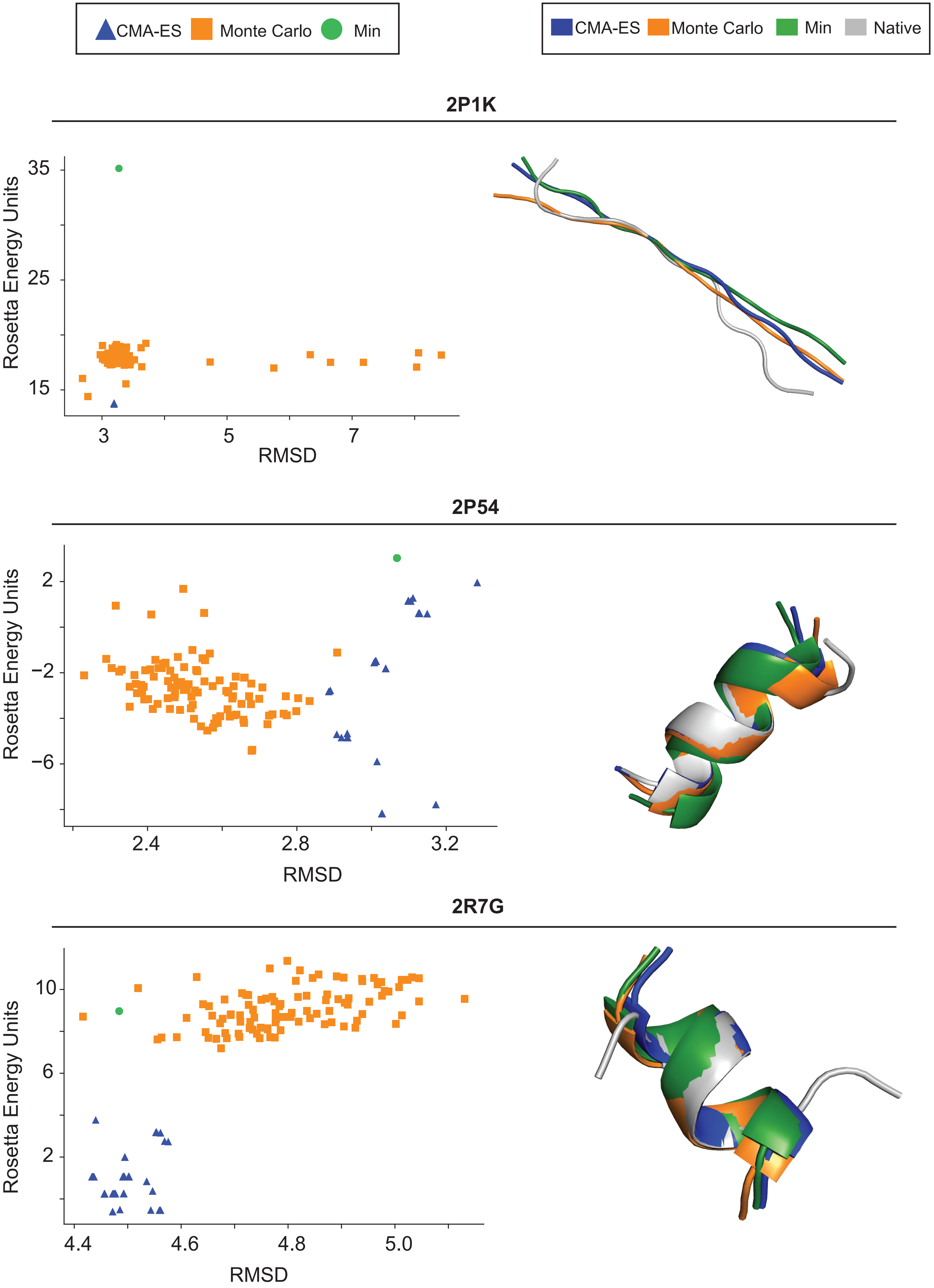

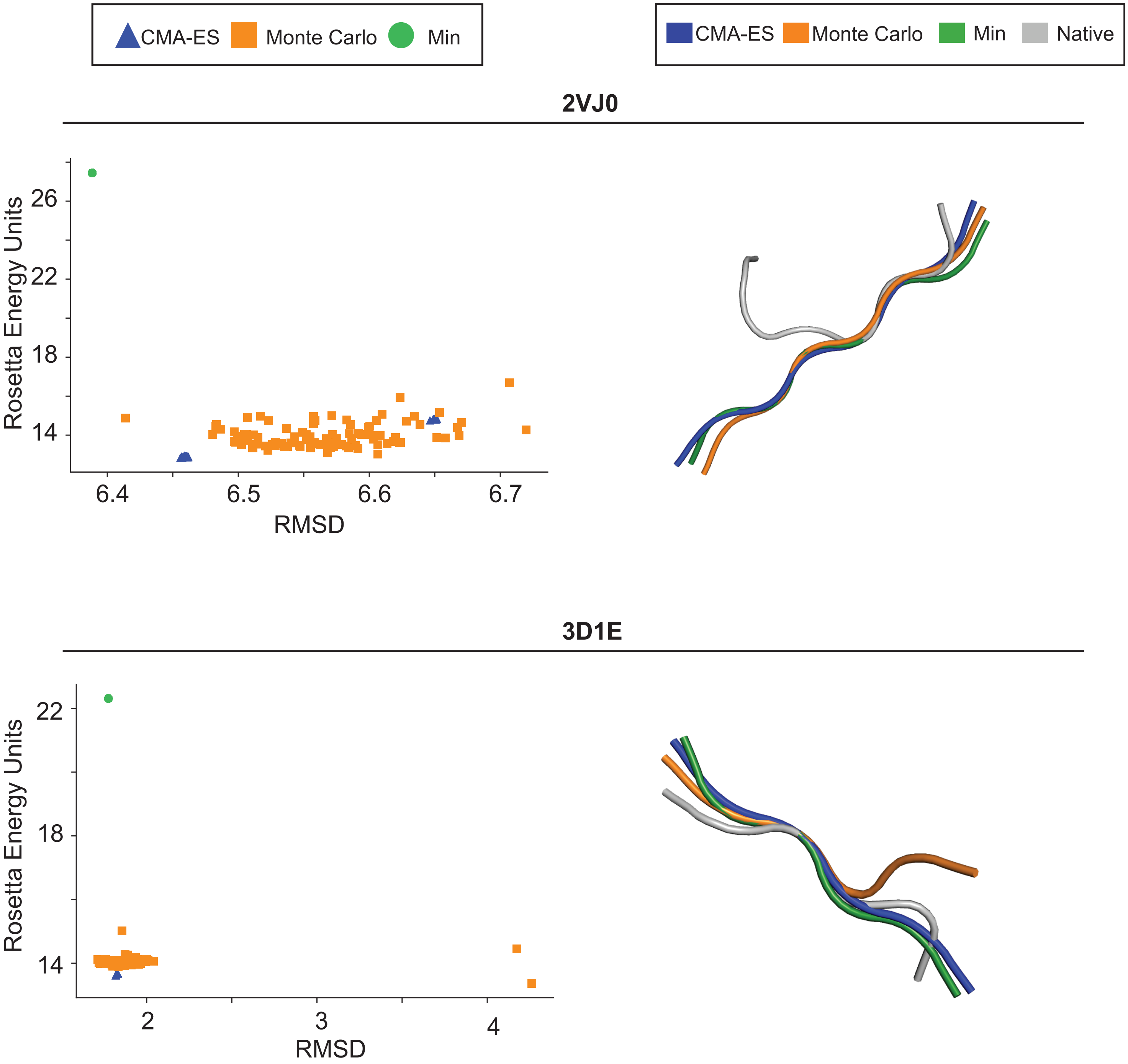

### Supplemental Figure 3

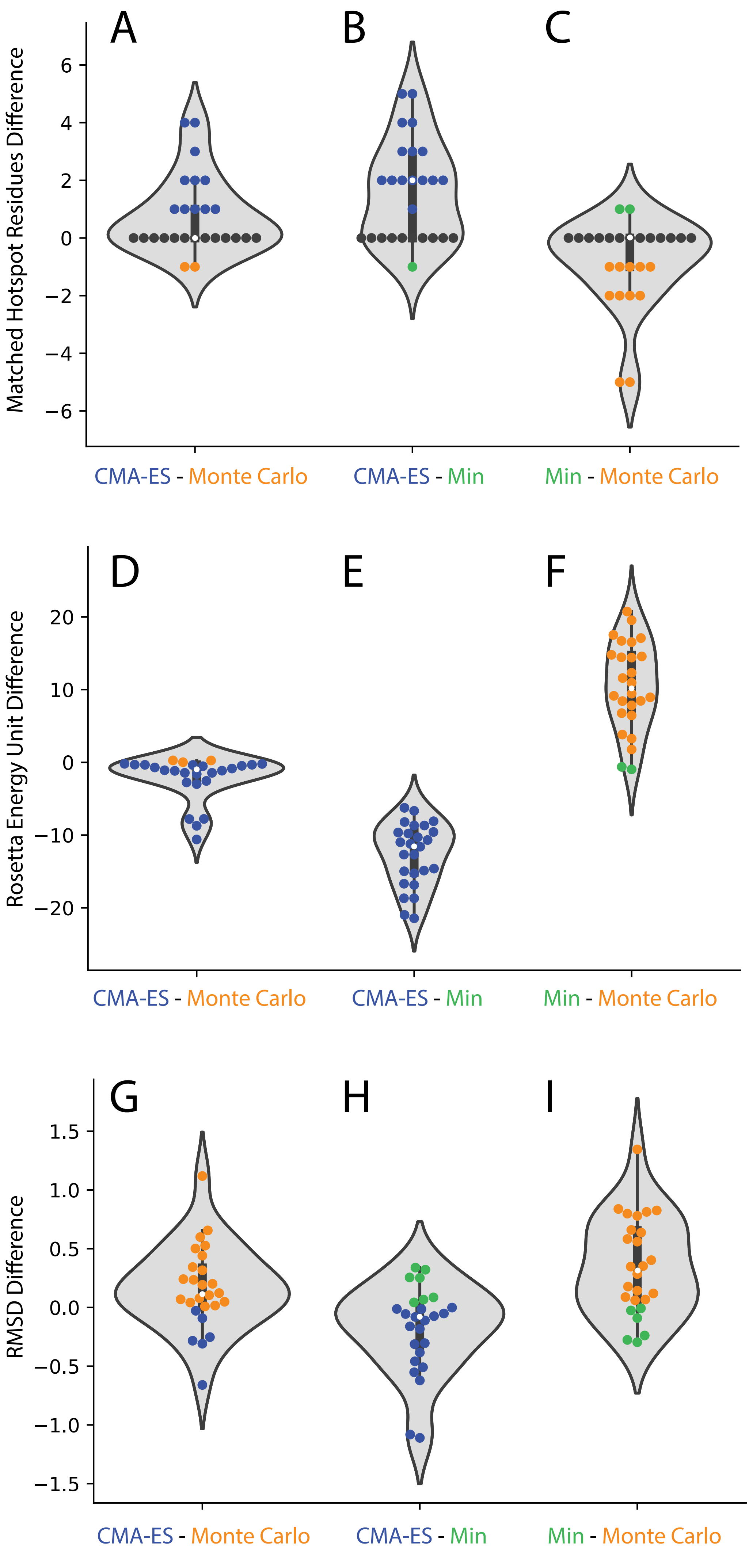
